# Pseudo-spectral angle mapping for automated pixel-level analysis of highly multiplexed tissue image data

**DOI:** 10.1101/2024.01.09.574920

**Authors:** Madeleine S. Durkee, Junting Ai, Gabriel Casella, Thao Cao, Anthony Chang, Ariel Halper-Stromberg, Bana Jabri, Marcus R. Clark, Maryellen L. Giger

## Abstract

The rapid development of highly multiplexed microscopy systems has enabled the study of cells embedded within their native tissue, which is providing exciting insights into the spatial features of human disease [1]. However, computational methods for analyzing these high-content images are still emerging, and there is a need for more robust and generalizable tools for evaluating the cellular constituents and underlying stroma captured by high-plex imaging [2]. To address this need, we have adapted spectral angle mapping – an algorithm used widely in hyperspectral image analysis – to compress the channel dimension of high-plex immunofluorescence images. As many high-plex immunofluorescence imaging experiments probe unique sets of protein markers, existing cell and pixel classification models do not typically generalize well. Pseudospectral angle mapping (pSAM) uses reference pseudospectra – or pixel vectors – to assign each pixel in an image a similarity score to several cell class reference vectors, which are defined by each unique staining panel. Here, we demonstrate that the class maps provided by pSAM can directly provide insight into the prevalence of each class defined by reference pseudospectra. In a dataset of high-plex images of colon biopsies from patients with gut autoimmune conditions, sixteen pSAM class representation maps were combined with instance segmentation of cells to provide cell class predictions. Finally, pSAM detected a diverse set of structure and immune cells when applied to a novel dataset of kidney biopsies imaged with a 43-marker panel. In summary, pSAM provides a powerful and readily generalizable method for evaluating high-plex immunofluorescence image data.

**Significance Statement:** Understanding the cellular constituents captured by highly multiplexed tissue imaging is a major limitation affecting the usability of these novel imaging methods. Many imaging experiments have uniquely designed staining panels, reducing the generalizability of cell classification models to new datasets. We present pseudospectral angle mapping (pSAM), which can compress high-dimensional image data into class representations. We demonstrate that the class representations generated by pSAM can be used to interpret high-plex image data and guide cell classification. Importantly, we also demonstrate that pSAM can generalize to new image datasets—collected with a different staining panel in samples from different tissues—without manual image annotation, subjective intensity gating, or re-training an algorithm.

## Introduction

The recent emergence of highly multiplexed tissue imaging modalities—systems that go beyond the traditional channel limit of immunofluorescence microscopy—has enabled the study of cells while maintaining spatial context, which is integral for furthering our understanding of immunity in human disease [1, 3-8]. Studying cells within native tissue is important for elucidating organizational principles of cells and studying cell:cell interactions that may otherwise be disrupted [7-11]. Tissue-destructive methods like single-cell RNA sequencing can provide specific information about gene expression in cells [12-14], but such methods lack the spatial context provided by imaging, and are therefore constrained in the information they can convey. Highly multiplexed imaging addresses the need for higher specificity in studying a variety of cell phenotypes in their native environment; multiple methodologies have been developed to image upwards of 40 protein markers in one field of view [3, 6, 15]. While these imaging methods are powerful and have the potential to unlock new insights into highly specific cell-cell interactions, there is still a need for robust and reliable methods for quantifying these types of images.

Most existing methods for characterizing high-plex images are focused on object-level analysis and annotation of cells [16, 17]. Specifically, cells are generally classified by their relative protein expression across several channels, either through a decision tree-based method or through clustering of mean pixel intensity (MPI) within each channel [16, 18, 19]. However, averaging intensity across a cell in an image loses potentially valuable organizational information stored in two-dimensional space such as signal localization within a cell. Additionally, immunofluorescence imaging captures only the two-dimensional section of cells and structures existing in a three-dimensional space. Many cells, particularly some populations of immune cells, can have protrusions that reach in and out of the imaging plane, potentially resulting in true detected signals not being assignable to a cell nucleus in the imaging plane [10, 20, 21]. Detection and characterization of cells in the imaging plane will provide integral information for spatial biology studies; however, the spatial context of the tissue is also worth understanding. We propose methods that can retain valuable information from the interstitial spaces between cells while providing an automated framework for rapid annotation of known cell types by collapsing the “spectral” or “channel” dimension of highly multiplexed images.

Annotation of cell classes with respect to cell type and cell state, such as activation state or proliferation state, in images is a particularly difficult computational task [10, 16, 22]. The boundary between cell class and cell state can be subjective, and ground truth is both noisy and difficult to acquire because human readers are not reliable in defining ground truth for cells in high-channel images [23]. Because of this scarcity of reliable ground truth and training data, methods like support vector machines or convolutional neural networks (CNNs) tend to perform poorly. Other methods classify cells by their protein expression (represented by MPI) across several markers, either through a decision tree-based method or through clustering of mean pixel intensities within each channel [16]. Cell typing in high-plex images is specific to the staining panel used in each imaging experiment, which means these models are often not generalizable to new datasets. For example, an imaging experiment which does not include CD4 in the staining panel cannot directly probe helper T cells, and models trained with this data would therefore lack the ability to classify helper T cells. Because of the variety of markers required to identify specific immune cell types and states, there is a need to develop more accessible and reliable methods for studying spatial immunity.

Here, we present an adaptation of a spectral angle mapping – a concept drawn from hyperspectral image analysis [24, 25]—to calculate pixel-level class representations by compressing the channel dimension of high-plex images through vector similarity calculations. While highly multiplexed immunofluorescence imaging does not physically collect hyperspectral data, parallels can be drawn between a single channel image of protein expression and a single wavelength band of a hyperspectral image. As this high-plex data is not truly hyperspectral or even spectral, we refer to data in the channel dimension of our images as “pseudospectral”. Reference vectors used for these vector similarity calculations can be generated for any arbitrary staining panel, making these methods more generalizable than current techniques, particularly across imaging experiments using different staining panels. We discuss methods for pre-processing image data and the generation of a library of “pseudospectra” for different immunofluorescence panels. We then demonstrate the efficacy of a pseudospectral angle mapping (pSAM) algorithm for condensing several channels of image data into class representation maps. Additionally, these class representations can aid in cell type annotation in highly multiplexed image datasets. These demonstrations were performed in two unique image datasets to show that the method can be applied across tissue types and staining panels. Overall, pSAM is rapid and adaptable, and will expedite the quantification of highly multiplexed imaging data for studying spatial immunity and spatial biology.

## Results

### Spectral angle mapping collapses high-dimensional image data at the pixel level

High-plex immunofluorescence imaging employs a staining panel of several protein markers which are carefully selected to probe cell populations and other molecules or structures of interest. Therefore, there is some a priori knowledge about which cell populations are captured by specific components of a staining panel. Pseudospectral angle mapping (pSAM) is a method for computing spatial likelihood maps for cell types probed by the markers used in a custom staining panel (Figure 1). In pSAM, a cosine similarity metric is used to evaluate how close a pixel vector is to a reference vector in an N-dimensional space, where N is the number of channels in an image. Given the wealth of literature defining immune cells based on their gene or protein expression [26-29], and the help of expert immunologists, we generated ideal pseudospectra as models of the cell types probed by two high-plex staining panels (Supplementary Figures 1-2). For example, an exhausted CD8 T cell will have high expression of the markers CD45, CD3, CD8, and PD1 on the surface of the cell, while a T regulatory cell will have high expression of CD45, CD3, and CD4 on the surface of the cell, and high expression of Foxp3 in the nucleus of the cell. Additionally, images were examined for cross-reactivity of the antibodies. For example, CD21 was selected to identify germinal center B cells in the kidney. However, this marker showed notably high signal in many cell nuclei within some tubules in the kidney. Therefore, we also generated a reference pseudospectrum for a “Tubule Nucleus” class, even though an antibody specific to tubule cell nuclei was not used in the staining panel. Representing each individual pixel in an image as a vector (or pseudospectrum), we then created low dimensional representations of highly multiplexed images called class maps. These class maps correspond to the likelihood of a given pixel belonging to a specific cell type.

**Figure 1.**
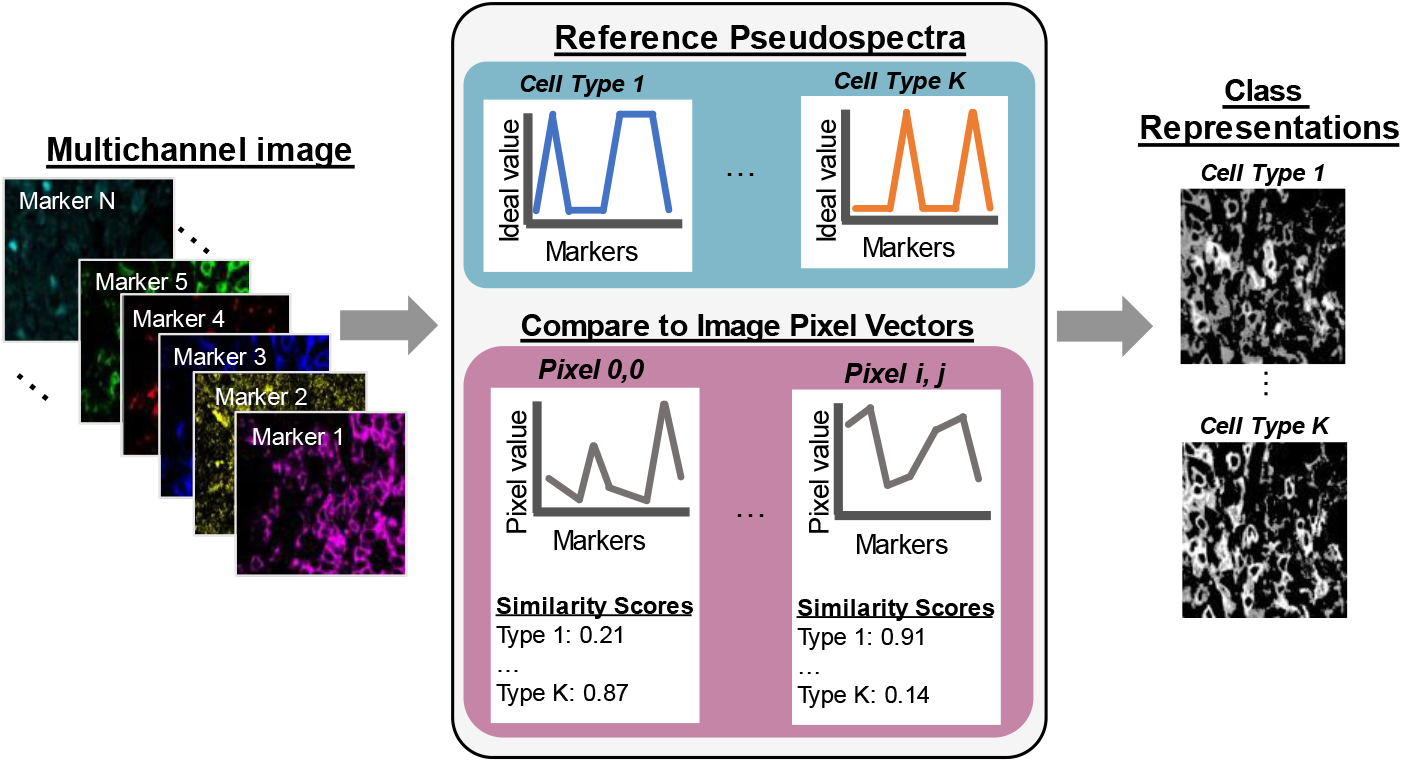
Pseudo-spectral angle mapping in highly multiplexed immunofluorescence images highlights pixels most similar to ideal reference profiles marker expression. Each pixel vector in a multichannel image, i.e., a pseudospectrum, is compared to ideal reference pseudospectra representative of each pre-determined class. The cosine similarity scores comparing the image pseudospectra to the references are mapped back to pixel coordinates to generate class maps of each cell type probed by the staining panel.

Ideal reference pseudospectra were defined by generating vectors of the expected relative protein expression by target cell populations. These expected expression profiles are compartment-dependent, meaning that certain pseudospectra occur in one area of a cell—for example the nucleus, while others occur in the cell cytoplasm or membrane. Reference pseudospectra were therefore defined by the location within a cell where a specific protein is expected to localize (Figure 2). For example, in a thirteen-marker panel used for staining colon biopsies from primary sclerosing cholangitis (PSC) patients, three markers (stains) probed cell nuclei, resulting in three unique ideal pseudospectra for the nucleus compartment (Figure 2A). The remaining ten markers are expected to be expressed on the cell membrane or within the cytoplasm of cells [30-32], collectively referred to here as the membrane compartment. Three of the reference pseudospectra for this compartment are shown in Figure 2B, with all reference profiles displayed in Supplementary Figure 1. In total, sixteen unique reference pseudospectra were generated from the thirteen-marker panel used in the PSC imaging experiment.

**Figure 2.**
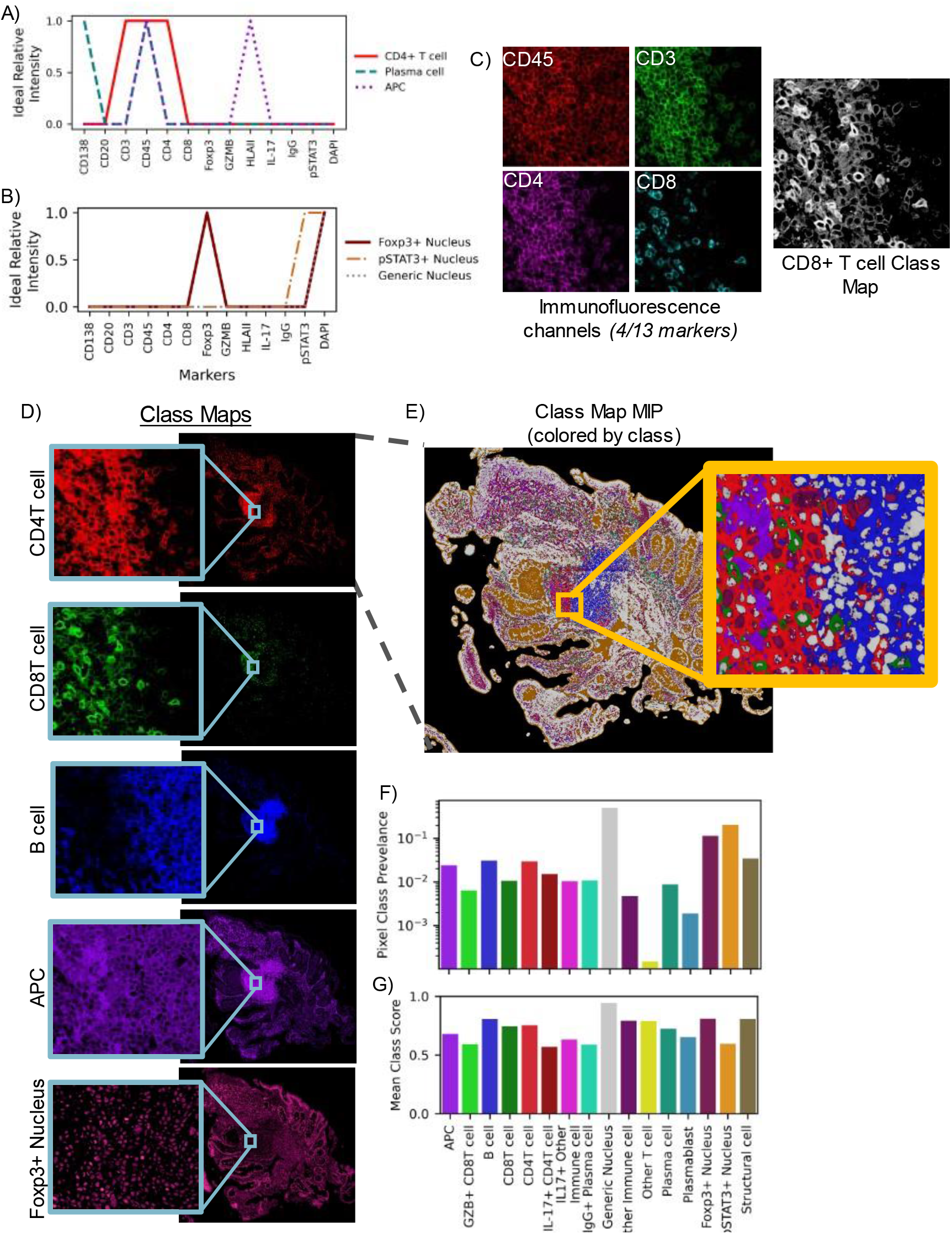
High-plex images are compressed into class representations maintaining full spatial scale using pseudo-spectral angle mapping (pSAM). Reference pseudospectra are designed for each imaging experiment. These references are specific to cell compartments such as the nucleus (A) or cell membrane (B), based on the molecular target of the marker. Aligned fluorescence images are used to generate similarity maps for each reference pseudospectrum (C). Four markers (CD3, CD45, CD4, CD8) are displayed in (C), but all thirteen channels are used to calculate the class maps. The pSAM class maps show the similarity of pixels to each cell type without the need to compare relative expression across multiple image channels in sample S1 (D). Given the maximum value at each x/y location, individual cells of different populations are clearly visible, even in densely packed areas (E). For the sample shown in (E), the prevalence of each pixel class within the tissue is calculated (F). The average cosine similarity score for each class of pixels in this example image is shown in (G). Prevalence and average score (F-G) are calculated over the pixels that fall within the tissue mask, while the slide background is filtered out as background (black in E).

Class similarity maps for a single reference pseudospectrum compress information from all markers in the panel. For a pixel vector to have a high similarity with the cytotoxic T cell class, that pixel must have high intensity values in the CD45, CD3, and CD8 channels and low intensity in all other channels, particularly CD4 (Figure 2C). Class maps of select reference pseudospectra for a full biopsy section are displayed in Figure 2D. A maximum intensity projection of all sixteen class maps (Figure 2E) revealed definitive structures—potentially even tertiary lymphoid structures, and object-level (or cell-level) variability into densely packed areas (Figure 2E, inset). Importantly, the class of the maximum score corresponded to the correct compartments, with nucleus-associated classes scoring highest in cell nuclei, and membrane-associated classes scoring highest in the pixels surrounding cell nuclei (likely membrane or cytoplasm). The class prevalence within this maximum intensity projection is displayed in Figure 2F. As expected, the highest prevalence class is the ‘generic nucleus’ class. Additionally, the ‘Other T cell’ class is also very low prevalence, which is expected, as T cell subsets that are negative for both CD4 and CD8 are generally rare. While very few pixels are most similar to the ‘Other T cell’ reference pseudospectrum, the average class score for this category was relatively high (Figure 2G). Conversely, the pSTAT3+ Nucleus class showed high prevalence and a relatively low average class score. Within the maximum intensity projection, the areas corresponding to the lumen of the intestine were associated with this class (Figure 2E, orange).

### Pixel-level class maps aid in cell phenotyping

By collapsing high-plex image data into pixel-level class representations, we are also able to infer likely cell phenotypes—merging pixel-level descriptors with object-level (cell-level) information. Class maps computed from pSAM were combined with instance segmentations of cell nuclei to assign cell classes (Figure 3). A cell’s overall class depends on the protein expression within both the nucleus compartment and membrane compartment. First, instance segmentations of cell nuclei were used to mask the nucleus-associated class maps and calculate overall representation scores for each reference pseudospectrum for each cell nucleus (Figure 3A, left). Similarly, dilated nucleus segmentations were used to mask the membrane-associated class maps to calculate scores for each membrane-associated class (Figure 3A, right). Each object was assigned a nucleus and a membrane phenotype according to the maximum class map score for each compartment. The predicted classes agreed well with the class representations and the fluorescence images themselves (Fig 2C&E, Figure 3B). These cell phenotype scores were validated by analyzing the MPI for each category of cells across all thirteen colon samples (Figure 3C). The intensity profiles for each detected cell phenotype corresponded well with the expected relative protein expression for the assigned cell type. This population-level validation demonstrates the potential of pixel-level analysis of highly multiplexed images to improve cell classification. Variation in cell class prevalence was seen across colon samples. Many samples had a plurality of IgG+ plasma cells, while others had a much smaller proportion of these cells (Figure 3D). Across all samples, a strong majority of cells were classified as having a ‘generic’ nucleus (neither Foxp3+ or pSTAT3+, Figure 3E). Foxp3+ nuclei were much more common than pSTAT3+ nuclei; however, it is worth noting that many samples showed markedly high Foxp3 signal in the villi of the intestine, which may have elevated this prevalence. In agreement with visual inspection of pSTAT3 images, many samples had no pSTAT3+ nuclei, and those that did had a very small proportion. As discussed above, a large number of pixels in the luminal space on the slide of S1 scored high for pSTAT3+ nucleus class (Figure 2E). Importantly, this did not matriculate through to cell class prediction. Note that we did not combine membrane and nucleus classes into canonical cell classes for this analysis (i.e. regulatory T cells would be CD4T cells or ICOS+ CD4T cells with a Foxp3+ nucleus). However, this may help to rapidly identify cross-reactivity of some markers if unexpected membrane+nucleus class combinations were to emerge.

**Figure 3.**
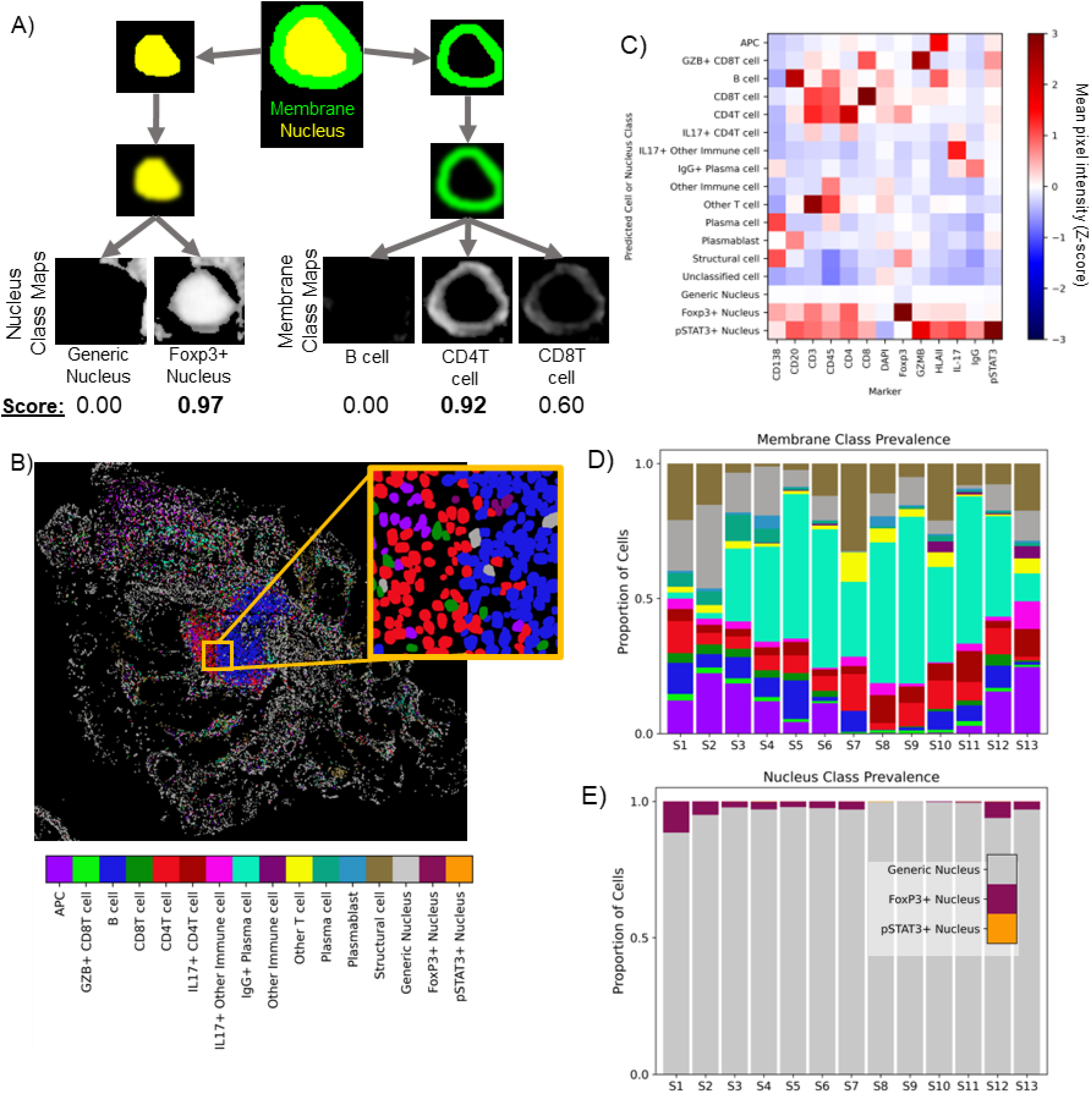
Class maps are combined with instance segmentations of cells to classify cells (A). Segmented nuclei and a dilated ring around each nucleus are used to mask class maps to generate class scores for each detected cell. The maximum score in each compartment is used to assign a compartment-specific class. Classifications based on cell membrane markers are shown for sample S1 in (B). Cell classes were validated using the mean pixel intensity (MPI) for all channels for each cell. MPI was z-scored across the full PSC dataset (thirteen biopsies, ∼885K cells), and the average Z-score for each channel is calculated for each cell class across all samples (C). Note that all cells receive a membrane class (top 13 rows) and a nucleus class (bottom 3 rows). Across the thirteen samples in the PSC dataset, pSAM detected varying distributions of cell classes in the membrane compartment (D) and the nucleus compartment (E). Notably, the generic nucleus class (gray) is a strong majority of cells, and pSTAT3+ nuclei (orange) are rare, but present and detectable in a few samples. The color bar in panel (B) is also applicable to panels (D) and (E).

### Generalizability to new immunofluorescence imaging experiments

High-plex immunofluorescence imaging experiments vary widely in the protein marker panel used to probe cells and tissue. Therefore, to evaluate the generalizability of pSAM, we applied the same method used on the 13-marker PSC dataset to a pilot set of three kidney samples imaged with the same 43-marker staining panel (Figure 4, Supplementary Table 2). Each kidney sample came from a patient with a different diagnosis: kidney transplant rejection, lupus nephritis, and angiomyolipoma. Samples from different pathologies were selected to demonstrate that pSAM is not only generalizable to new staining panels but is also functional across different pathologies imaged with the same markers, without the need for computational batch correction. From the 43-marker panel, we defined 38 unique reference pseudospectra using 30 of the 43 markers. 34 of these references were membrane-associated pseudospectra and four were nucleus-associated pseudospectra (Supplementary Figure 2). The higher plex staining panel used in the kidney dataset allowed for more than double the number of cell types and states to be detected in this dataset relative to the 13-plex PSC dataset. In each sample, despite coming from patients with different diagnoses, areas of inflammation and areas with structural cells qualitatively agree with the raw images. Additionally, pixel class prevalence trends as expected, as nucleus-associated classes had the highest prevalence, and more inflamed samples showed a higher prevalence of pixels showing high similarity to immune cell reference pseudospectra (Supplementary Figure 4). Specifically, the kidney transplant sample has by far the highest prevalence of lymphocyte-associate pixels (T and B cells), and this sample has three large clusters of lymphocytes visible in the image (Figure 4A). Cell classes were also computed using the instance segmentation masks and the pixel class representations (Figure 4A). Validation with MPI across each predicted class of cells showed that each cell phenotype had the expected relative protein expression (Figure 4B). The trends for protein expression of a given cell type are similar across these samples from different pathological conditions (Supplementary Figure 4). All 43 markers were used for MPI validation, while only 30 markers were used for cell classification. Therefore, a few surprising trends were found with the remaining 13 markers. For example, ROR-γt (not used in cell classification) was relatively high in SLAMF7+ plasma cells. Further investigation is needed to determine whether this finding is associated with cross-reactivity of antibodies or spatial overlap of signal from neighboring cells. Further validation of the pSAM cell classification showed that the prevalence of immune cell populations is highest in the transplant rejection sample, which corresponds with visual analysis of the full biopsy sections (Figure 4C). Additionally, the generic nucleus class is by far the most abundant across samples, as expected (Figure 4D).

**Figure 4.**
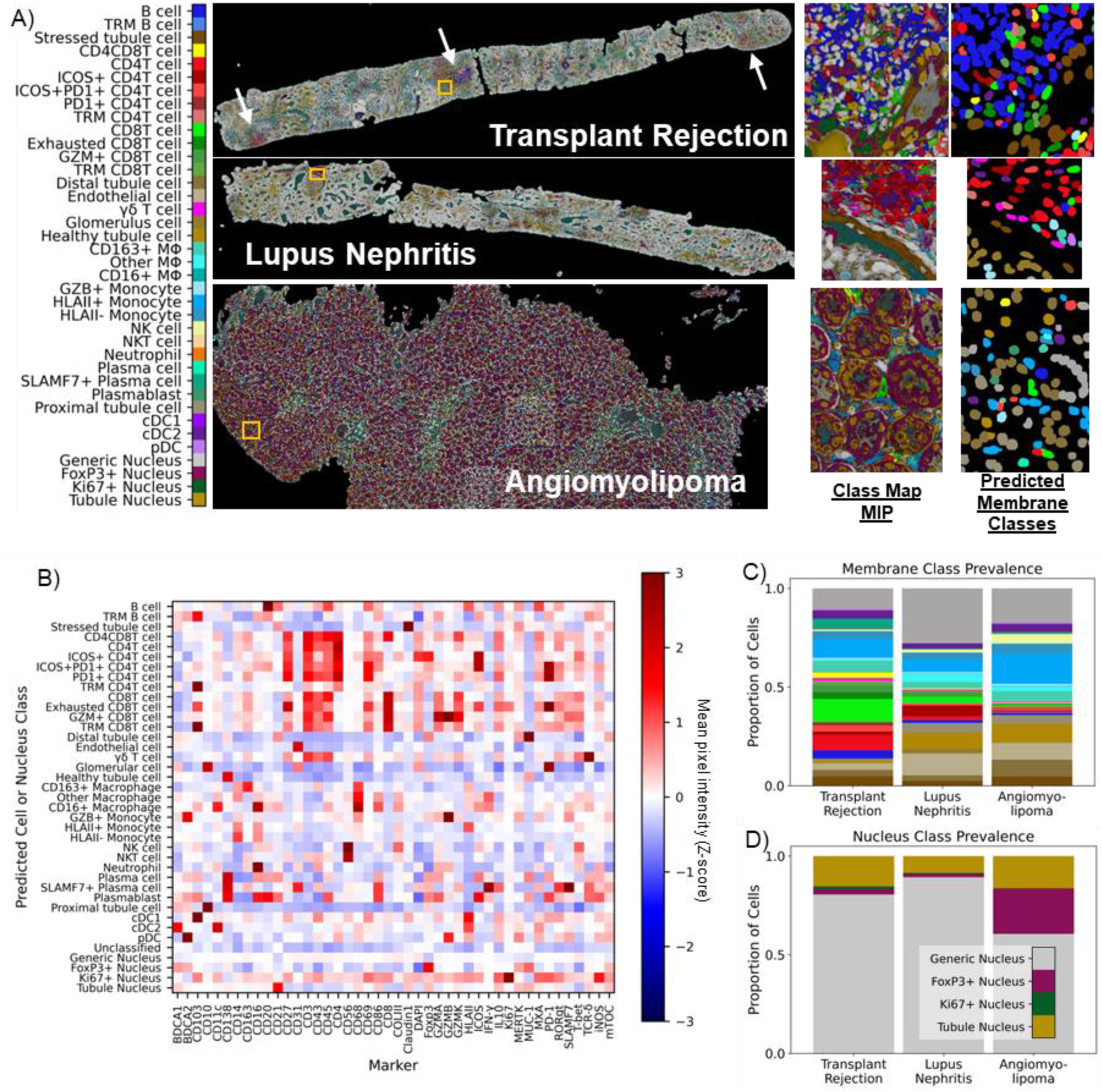
pSAM is readily generalizable to new datasets. A) pSAM was run on three kidney samples from a dataset stained with a 43-marker panel, each coming from a patient with a unique diagnosis. pSAM was used for cell classification in these three samples, with each cell receiving a class assignment for one of 35 membrane classes and one of four nucleus classes. White arrows in the transplant rejection image point to large clusters of lymphocytes. B) The z-score of the population mean pixel intensity (MPI) for all stains in the 43-marker panel is compared across all cell membrane and nucleus classes. Note that all cells have both a membrane class and a nucleus class. C) A comparison of the relative populations in each sample shows that the immune populations detected shows a diverse and abundant population of immune cells, corresponding with visual inspection of the images. Similarly, tubule and generic nuclei, (gold and gray, respectively) are by far the most abundant nuclei in all three samples (D). Notably, Foxp3 shows cross-reactivity in many tubules in the kidney particularly in the angiomyolipoma sample (A, class map). This is reflected in the cell classification with an overabundance of Foxp3+ nuclei detected. For clarity, cell classes are color by a broader umbrella depending on their lineage (structural cells in brown, B cells in blue, CD4 T cells in red, etc.). The color bar in (A) is also applicable for panels (C) and (D).

## Discussion

Pseudo-spectral angle mapping (pSAM) provides a new tool for analyzing pixels and cells in highly multiplexed microscopy images. Compressing multi-channel image data into class representations can help to facilitate rapid viewing of these complex images and evaluate the cellular constituents of a biopsy sample. In this work, we demonstrated the benefits of implementing pSAM for pixel-level and cell-level classification in two different image datasets imaged with very different staining panels. First, in thirteen-plex images of colon biopsies from PSC/IBD patients, pSAM class maps demonstrated that condensing the channel dimension of these images allows for rapid visualization of unique cell types in dense regions of immune infiltrates. Not only were these class maps valuable at the pixel and regional level, combining them with a basic cell-by-cell masking algorithm produced sensible cell classifications. Cell class annotations in high-plex images are particularly difficult for human readers. Since human readers are the “gold standard” for this task, and they are noisy and inconsistent [23, 33, 34], supervised image classification algorithms using image annotations are not ideal for this task. pSAM provides a means for classifying cells that does not rely on manual image annotations, and therefore does not depend on noisy and unreliable truth that is difficult and costly to obtain. In the 43-plex kidney images, pSAM detected a wide range of cell classes with highly variable prevalences, suggesting that it can accurately detect rare cell types. However, there were some discrepancies in the angiomyolipoma sample with tubule cells often classified as monocytes and plasmacytoid dendritic cells (pDCs). Upon analysis of the images the tubules in this sample are highly irregular and have high levels of expression of CD14 (a marker for myeloid cells) and BDCA2 (a marker for pDCs). Additionally, the markers in the staining panel that are intended for tubules stain the brush border of the tubule structure, and not the cell membrane or the cytoplasm near the nucleus (Supplementary Figure 5). The localization of the tubule marker signal to the tubule structure and not the individual cells would result in poor cell classification when classifying a cell based on the nucleus dilation methods shown here. However, the class maps show high agreement with the tubule classes at the brush border of the tubule and therefore maintain important contextual information that would be lost in a cell-centric classification method. Therefore, pSAM provides a rapidly adaptable tool for assessing pixel classes which are helpful for inferring cell type prevalence in images of diverse populations of cells.

High-plex imaging is often used for biological discovery, and the stains used are subject to change from experiment to experiment as researchers toggle the cell types and populations they want to probe. The acquisition of robust and reliable ground truth for every new imaging experiment is costly and time-consuming, even if the quantity of manual annotations is reduced by fine-tuning an existing model. Supervised classification of cells in highly multiplexed images can be particularly difficult because image annotations are both difficult to acquire and specific to the staining panel used for the imaging experiment [34]. Therefore, models trained for cell classification in high-plex images are not typically generalizable to new datasets [35]. Additionally, identification of rare cell types is also problematic because they or often not represented in training data. pSAM requires no image annotations – no noisy ground truth from human readers. While this method can be interpreted as “supervised”, all supervision comes from reference pseudospectra generated from the staining panel. Reference pseudospectra can be easily generated for each unique imaging experiment, making pSAM a desirable method for classifying pixels and cells in high-plex images. Additionally, while not demonstrated in this work, vector similarity metrics can also be used for spectral unmixing [36]. In densely packed regions of cells, different cell types can have overlapping signal at the cell membrane [37]. This overlapping signal problem, or ‘mixed signal’ can also potentially be addressed through spectral angle methods by allowing linear combinations of reference pseudospectra [24].

Despite the massive benefits of using pSAM for pixel and cell classification, the method is not without its limitations. Pixel-level analyses require pixel-level calculations. Although a cosine similarity function is very quick to calculate, whole-slide images tend to be several giga-pixels in size, so pSAM is a compute-heavy method, even with optimal parallelization. Also, the argument could be made that interstitial pixels not associated with cells could simply contain noise or nonspecific antibody staining. However, if these pixels have a very high similarity to a reference pseudospectrum, it is much less likely to be non-specific staining, as multiple markers would have preferentially stained that location. High-plex imaging experiments can also be used to probe unexpected cell types or characterize the interstitial or acellular spaces in tissue [38]. The pSAM algorithm presented here can only detect the classes defined by the reference pseudospectra. In general, staining panels (even high-plex panels) are experimentally designed to probe known cell types or states, so the ‘you can only find what you look for’ limitation of pSAM may not be a limitation for many imaging experiments, particularly those in which researchers are interested in the spatial distributions of known or specific cell types. Liu et. al. has developed another pixel-level analysis method and demonstrated that random sampling of pixels yields reproducible classes of pixels [3]. While powerful, this sampling could miss rare cell types that could be easily detected with pSAM. Finally, pSAM cell classification is optimized to classify cells with small cell bodies – including lymphoid cells, endothelial cells, and some myeloid cells. As mentioned earlier, tubule cells in the kidney are often characterized by signal at the brush border of the tubule, which can be far away from the tubule cell nucleus. A simple nuclear dilation might not catch the associated cell signal. In this work, we describe pSAM cell classification using a dilated nucleus segmentation to classify the cell. More advanced whole-cell segmentation would improve pSAM cell classification.

Overall, pSAM is a simple and elegant solution for classifying pixels and cells in highly multiplexed microscopy data. While the method has its limitations, it provides a generalizable and easily implementable solution for compressing high-channel image data into lower dimensional representations. Importantly, pSAM mitigates the need to collect manual annotations on images or individual cells while probing the cell classes and states that a high-plex panel was designed to detect.

## Materials and Methods

### Sample and image acquisition

Thirteen colon biopsy samples from patients diagnosed with primary sclerosing cholangitis (PSC) and/or inflammatory bowel disease (IBD) were acquired from the Human Tissue Resource Center at the University of Chicago (HTRC). Additionally, three kidney samples were selected from a second dataset acquired from the HTRC. One kidney sample was from a patient diagnosed with lupus nephritis, one from a patient with mixed rejection of renal allograft (referred to later as transplant rejection), and one from a patient with angiomyolipoma.

A 5 µm section of each biopsy was mounted on a functionalized coverslip for iterative staining and imaging. Colon samples were iteratively stained and imaged with a thirteen-marker panel (Supplementary Table 1) and kidney samples were iteratively stained and imaged with a 43-marker panel (Supplementary Table 2). The PhenoCycler protocol was used for iterative staining [39]. All samples were imaged on an Andor Dragonfly spinning disk confocal microscope with a 40x objective lens. For colon samples, each cycle of imaging included a nucleus stain (DAPI) imaged at 405nm and three other stains from the panel imaged at 488nm, 561nm, and 637nm. For kidney samples, each imaging cycle contained DAPI and four other stains, with an additional channel at 730 nm. Blank imaging cycles (no fluorescence reporters except for DAPI) were acquired before and after all staining cycles to capture background tissue autofluorescence. Resulting full-section images had a 0.1507 µm pixel size.

### Image preprocessing

Images were acquired in 2048 x 2048 pixel fields of view with a 205 pixel (∼10%) overlap with neighboring tiles. Ashlar stitching and alignment software [40] was used to stitch all image tiles into full-section composites and align all imaging cycles to the DAPI channel from the first staining cycle [40]. Ashlar performance was visually checked across all samples. After aligning all image channels, the first blank cycle of imaging was used for background subtraction and spectral normalization of all stained images. First, each channel of the blank cycle was subtracted from the corresponding fluorescence channel in all imaging cycles. Each imaging wavelength has a different dynamic range, so the subtracted images were also divided by the standard deviation of the background image to standardize dynamic range across imaging wavelengths. After standardization relative to imaging wavelength, images were min-max normalized to the 99^th^ percentile.

### Instance segmentation of cells

The DAPI channel of each image was used to generate instance segmentations of all cells in each full-section image. The ‘nuclei’ model in Cellpose2.0 [41] was fine-tuned for each dataset to predict individual cell masks. In a test set of 50 image tiles per dataset (512 x 512 pixels), this fine-tuned model achieved an F1-score of 0.83 ± 0.13 for the PSC dataset, 0.86 ± 0.11 for the Lupus dataset, and 0.70 ± 0.16 for the transplant dataset. These test sets were comprised of images from five separate biopsies for each dataset. (Note that the three kidney samples used for the generalization dataset were selected from a larger kidney database). The model tuned for the lupus dataset was also used to segment cell nuclei for the angiomyolipoma sample. Nucleus segmentations were dilated by ∼1 µm (7 pixels) to achieve approximate whole-cell segmentations. Voronoi tessellation was used to avoid overlap of expanded nuclei in areas of crowded cells. Cell membrane segmentations were generated by subtracting each nucleus segmentation from the corresponding whole-cell segmentations (Figure 3A).

### Defining reference pseudospectra

The specific pseudospectra were generated for each panel were generated through literature review and with expert immunologist input. Because certain markers are known to localize to different compartments of the cell (i.e. nucleus, cytoplasm, or membrane), compartment-specific pseudospectra were defined. All markers from the staining panel were included in each pseudospectra (Supplementary Figure 1). For the PSC dataset, these reference (or ‘ideal’) pseudospectra were defined with binary values (0 or 1) for each marker. In more complex panels such as the kidney panel, there are situations in which a marker is expected to be expressed at multiple levels (high/low/no expression). In these cases, a value of 0.5 was used to capture low expression (Supplementary Figure 2). Importantly, marker order was held consistent across all defined references. The tubule nucleus reference spectra used for the kidney data defined CD21 as high in the tubule nucleus compartment. While this is not necessarily expected biologically, there was significant cross-reactivity of the CD21 marker in tubule nuclei, particularly in pathological states.

### Pseudospectral angle mapping (pSAM)

All image channels were consistently stacked in the order defined by the reference pseudospectra. The images in the stack were previously min-max normalized to the 99^th^ percentile and stored as 8-bit images (255 max pixel value). Prior to pSAM calculations, each channel was one-hot encoded to match the range of the reference pseudospectra (0-1). A pixel vector (pseudospectrum) extracted from the image data, was defined as the value of all markers at a single (x, y) position. Each pixel vector in the image was compared to all reference pseudospectra by computing the cosine similarity between the two vectors (Eq. 1, range 0-1). For visualization, the class maps were multiplied by a mask of the whole tissue, bringing the coverslip background to zero. These tissue segmentations were generated by gaussian filtering (sigma = 75) and thresholding the DAPI channel of the image stack.

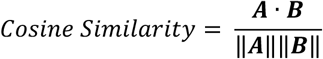

### Cell phenotyping

The PSC dataset included thirteen protein markers. Sixteen reference pseudospectra were generated from this thirteen-marker panel including thirteen unique cell membrane pseudospectra and three unique cell nucleus pseudospectra. Notably, there are more cell classes that could exist in these inflamed colon samples which cannot be captured by this thirteen-marker staining panel. We therefore expected a portion of the detected cells in each sample to remain unclassified. The kidney dataset contained 43 markers (30 used for pSAM), for which 42 unique reference pseudospectra were generated: 34 membrane-associated pseudospectra and four nucleus-associated pseudospectra. For the kidney dataset, we also expect a contingent of unclassified cells; some that are not captured by the staining panel, and some that are not captured by this set of reference pseudospectra. We generated 38 total reference spectra fro this panel, but more tractable reference spectra could be generated from these 43 markers.

To optimize our cell phenotyping, a threshold of 0.5 was applied to all class maps to reject low cosine similarity scores. Cell membrane segmentations were computed by subtracting the nucleus segmentation from the whole-cell segmentation. For each cell predicted by the fine-tuned Cellpose model, we computed a score for all computed class maps using the following protocol (Figure 3A). A “soft” mask of each nucleus segmentation was created using a distance transform to blur the edges of the binary mask. These “soft masks” were multiplied by all class maps associated with the nucleus compartment. The membrane mask (dilated nucleus segmentation mask – nucleus segmentation mask) was also blurred using a distance transform and multiplied with all class maps associated with the membrane compartment. The mean score of the masked class maps was computed for all reference pseudospectra for each cell. Each cell was given a nucleus classification and a membrane classification based on the maximum score for each compartment. A cell received a membrane class of “unclassified” if it scored zero for all membrane classes. If the maximum score for the nucleus compartment was zero, the cell received a nucleus classification of ‘generic nucleus’.

### Cell class validation

Cell classes were validated with the MPI in each cell segmentation. Briefly, the MPI was calculated for all cell segmentations in all image channels, even those not used for pSAM calculations. The MPI was z-scored across the full dataset: all thirteen samples for the PSC dataset, and all three kidney samples for the generalization experiment. The average z-score for each class is displayed for all image channels in heatmaps in Figures 3C, 4B, and Supplementary Figure 2.

## Acknowledgments

M.S.D. is grateful for the support of the Eric & Wendy Schmidt AI in Science Postdoctoral Fellowship, a Schmidt Futures program. This research was supported by the National Institute of Allergy and Infectious Diseases (NIH) under award numbers U19 AI082724 (M.R.C.), R01 AR055646 (M.R.C.), R01 AI148705 (M.R.C.), and U01 CA195564 (M.L.G.). The content is the responsibility of the authors and does not necessarily represent the official views of the NIH. Funding was also provided by the Department of Defense, award number LR180083 (M.R.C), and the GI Research Foundation of the University of Chicago Medicine Digestive Diseases Center (M.S.D). Computational resources and support were provided by NIH S10 OD025081 Shared Instrument Grant (M.L.G). Special thanks to Chun-Wai Chan, MSc for computational support and guidance, and to Cezary Ciszewski of The University of Chicago Human Disease and Immune Discovery for help with image acquisition.

