## Supplemental Figures and Tables for "Pseudo-spectral angle mapping for automated pixel-level analysis of highly multiplexed tissue image data"

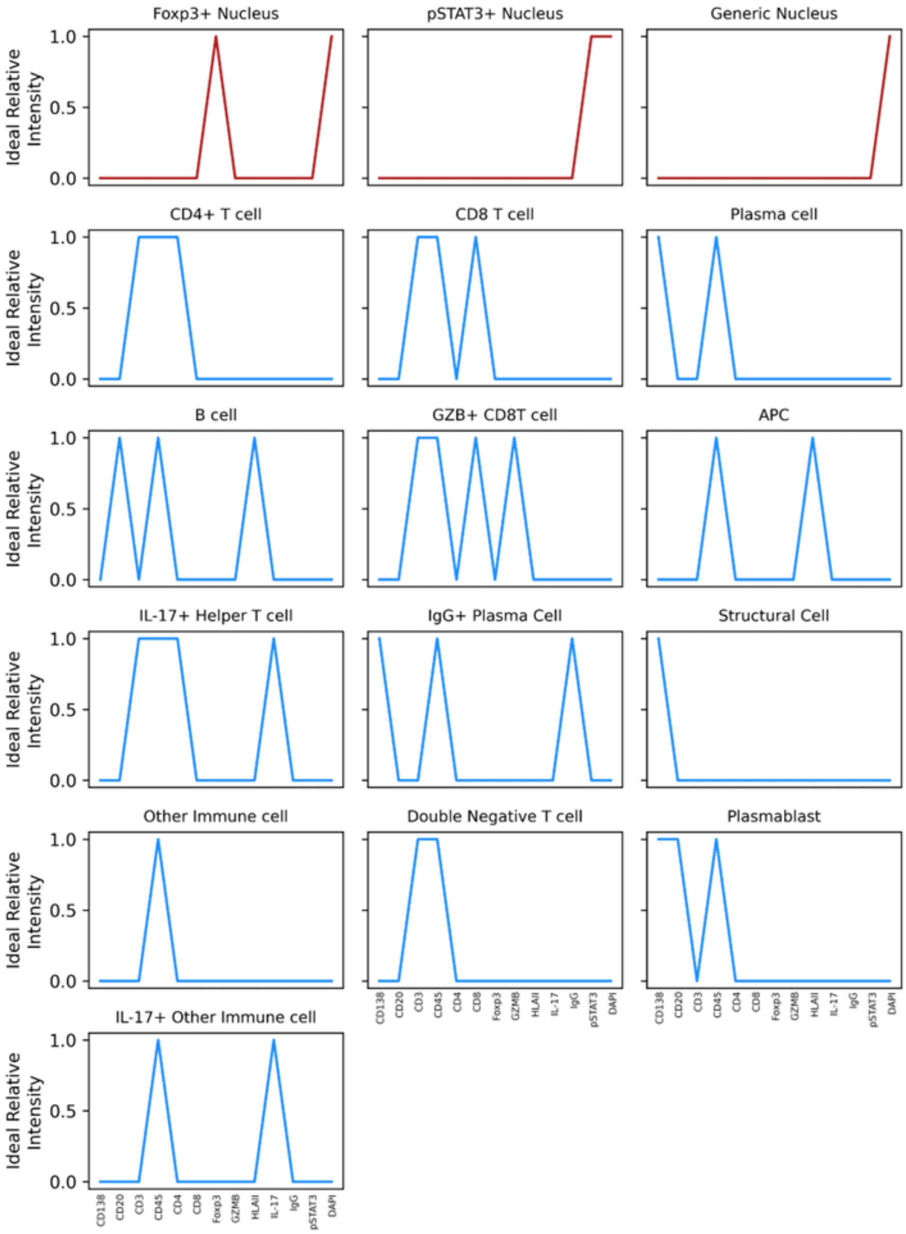


**Supplementary Figure 1.** All reference pseudospectra are displayed for the thirteen-marker panel used to image the PSC/IBD dataset. Pseudospectra designed to be high in the cell nucleus are displayed in red, while pseudospectra designed to be high in the cell membrane are displayed in blue.

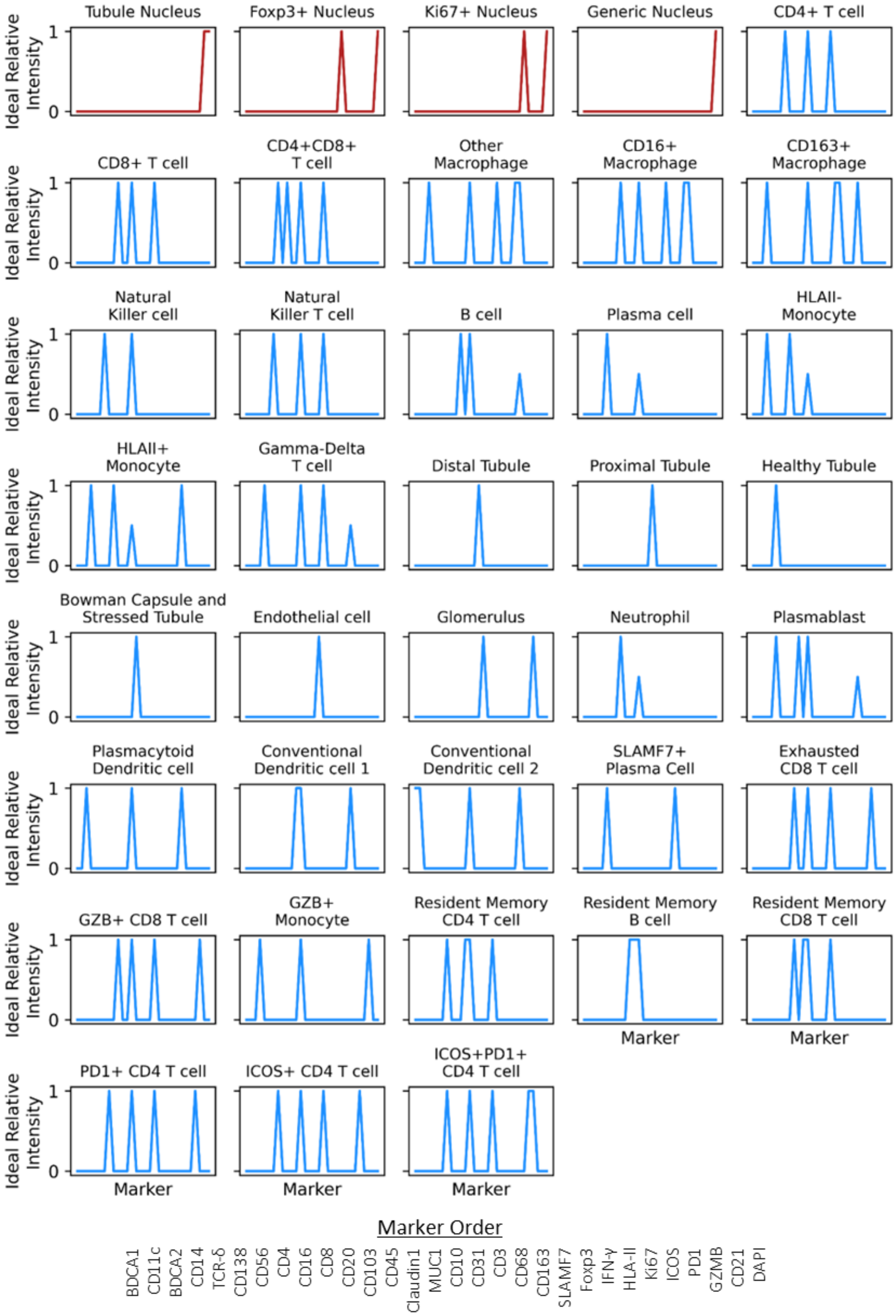


**Supplementary Figure 2.** All reference pseudospectra are displayed for the 30-marker panel used to image the kidney dataset. Pseudospectra designed to be high in the cell nucleus are displayed in red, while pseudospectra designed to be high in the cell membrane are displayed in blue. The marker order (held consistent across all x-axes in this figure), is displayed at the bottom of the figure.

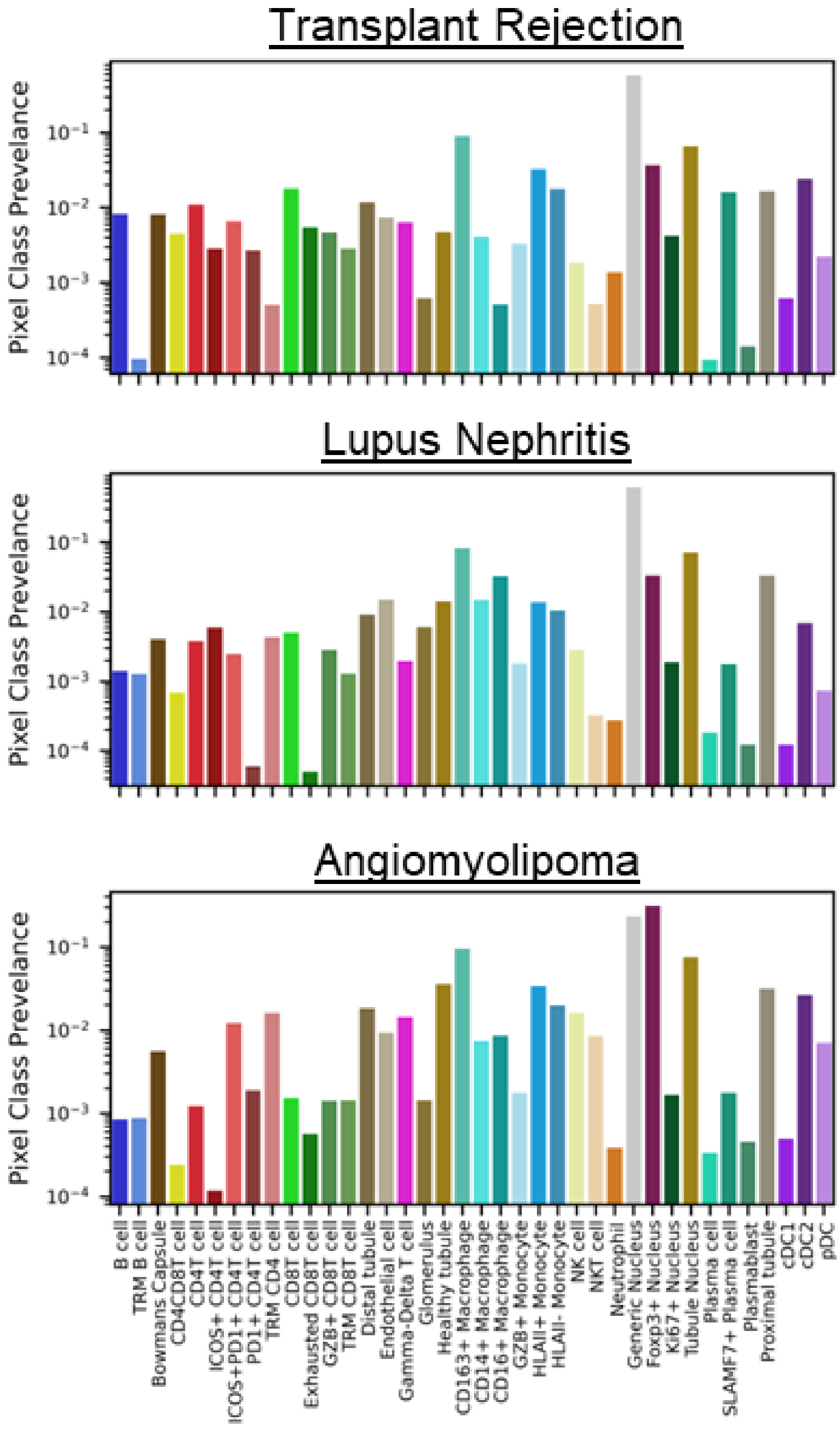


**Supplementary Figure 3.** Pixel class prevalence (log scale) for all classes is displayed for all three kidney biopsies: transplant rejection (top), lupus nephritis (middle), and angiomyolipoma (bottom).

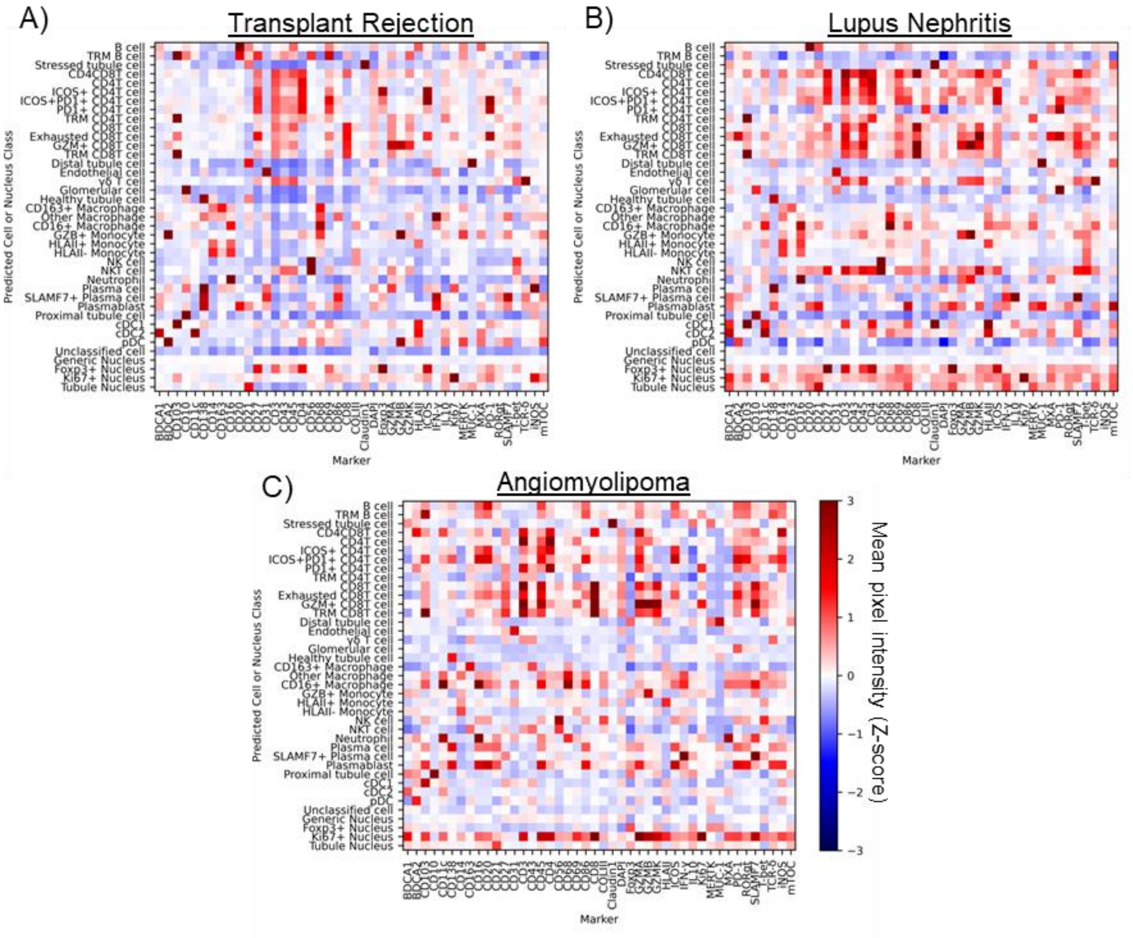


**Supplementary Figure 4.** Mean pixel intensity (MPI) for all channels in the 43-marker kidney panel by cell class for kidney transplant rejection (A), lupus nephritis (B), and angiolipoma (C). MPIs were z-scored across each channel for each sample.

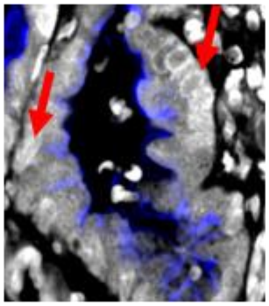


**Supplementary Figure 5.** Representative CD10 expression (distal tubule marker) in a kidney sample. CD10 signal resides outside of the range of nucleus dilation for many tubule cells (red arrows).

| Marker | Target | Vendor | Cat. No. |
|---|---|---|---|
| CD3 | Pan T cell marker | abcam | ab205228 |
| Foxp3 | Treg nuclei | eBioscience | 14-4776-82 |
| HLAII | Antigen presentation | Abcam | ab7856 |
| CD4 | Helper T cells | abcam | ab181724 |
| CD20 | B cells | abcam | ab236434 |
| IL-17 | TH17 cells | Novus Biologicals | AF-317-NA |
| CD8 | Cytotoxic T cells | Invitrogen | MA5-13473 |
| CD138 | Plasma cells/Villi | ThermoFisher Scientific | MA1-10091 |
| pSTAT3 | Carcinogenic/Dysplastic cells | abcam | ab219593 |
| GZMB | Activated CD8 T cells | cell signaling | 79003SF |
| IgG | Activated Plasma Cells | abcam | ab226069 |

**Supplementary Table 1.** All antibodies used in the PSC/IBD staining panel, including the expected expression for each marker.

| Marker | Target | Vendor | Cat. No. |
|---|---|---|---|
| CD31 | Endothelial cells | abcam | ab226157 |
| T-bet | TH1 nuclei | Cell signaling | 27112SF |
| CD103 | Resident memory cells/cDCs | abcam | ab254201-1001 |
| MUC-1 | Distal tubules | Novus Biologicals | NBP2-44658 |
| Foxp3 | Treg nuclei | eBioscience | 14-4776-82 |
| BDCA1 | cDCs | Novus Biologicals | NBP2-70345 |
| CD20 | B cells | Invitrogen | 14-0202-82 |
| CD10 | Proximal Tubules | Abcam | ab256494 |
| MXA | Interferon Response | R&D | AF7946 |
| GZMB | Activated CD8 T cells | cell signaling | 79003SF |
| CD68 | Macrophages | eBioscience | 14-0688-82 |
| BDCA2 | pDCs | Novus Biologicals | AF1376 |
| CD27 | B memory cells | abcam | ab272072 |
| GZMA | Activated CD8 T cells | abcam | ab251499 |
| PD-1 | CD8 T exhaustion and TFH cells | Cell signaling | 63815SF |
| CD86 | T cell activation | Cell signaling | 76755 |
| CD45 | Pan-immune cell marker | Biolegend | 11-9459-42 |
| ICOS | TFH cells | Cell signaling | 39740SF |
| GZMK | Activated CD8 T cells | abcam | EPR24601-164 |
| CD4 | Helper T cells | abcam | ab181724 |
| RORγt | TH17 nuclei | ThermoFisher Scientific | PA534164 |
| mTOC | Cell:cell interaction (with CD43) | abcam | ab27074 |
| CD43 | Cell:cell interaction (with mTOC) | Novus Biologicals | NBP2-34775 |
| CD69 | Lymphocyte activation | abcam | ab234512 |
| IFN-γ | Macrophages | abcam | ab231301 |
| CD163 | M2 macrophages | Novus Biologicals | NBP110-40686 |
| CD21 | Germinal center B cells* | abcam | ab271855 |
| HLAII-DR | Antigen presentation | abcam | ab7856 |
| CD8 | Cytotoxic T cells | Invitrogen | MA5-13473 |
| CD16 | Neutrophils, NKs, Monocytes, Macrophages | Cellsignaling | 72204SF |
| TCR-δ | γδ T cells | Santa Cruz | sc-100289 |
| CD138 | Plasma cells | ThermoFisher Scientific | MA1-10091 |
| CD11c | cDCs | abcam | EP1347Y |
| CD3 | Pan T cell marker | abcam | ab271850 |
| IL-10 | TH1/TH2 cells | Novus Biologicals | AF-217-NA |
| Ki67 | Cell proliferation | abcam | ab279657 |
| Claudin1 | Bowman's Capsule** | abcam | EPR121871 |
| COLIII | Collagen | Proteintech | 22734-1-AP |
| MerTK | Phagocytosis | abcam | ab271851 |
| iNOS | Inflammatory macrophages | abcam | ab239990 |
| CD14 | Monocytes, macrophages | abcam | ab230903-1001 |
| SLAMF7 | Activated Plasma cells | Cell signaling | 98611S |
| CD56 | NK and NKT cells | OriGene | CF506208 |

**Supplementary Table 2.** All antibodies used in the kidney staining panel, including the expected expression for each marker.
